# Local Contractions Regulate E-Cadherin Adhesions, Rigidity Sensing and Epithelial Cell Sorting

**DOI:** 10.1101/318642

**Authors:** Yian Yang, Emmanuelle Nguyen, Gautham Hari Narayana Sankara Narayana, Melina Heuzé, René-Marc Mège, Benoit Ladoux, Michael P. Sheetz

## Abstract

E-cadherin is a major cell-cell adhesion molecule involved in mechanotransduction at cell-cell contacts in tissues. Since epithelial cells respond to rigidity and tension in the tissue through E-cadherin, there must be active processes that test and respond to the mechanical properties of these adhesive contacts. Using sub-micrometer, E-cadherin-coated PDMS pillars, we find that cells generate local contractions between E-cadherin adhesions and pull to a constant distance for a constant duration, irrespective of pillar rigidity. These cadherin contractions require non-muscle myosin IIB, tropomyosin 2.1, α-catenin and binding of vinculin to α-catenin; plus, they are correlated with rigidity-dependent cell spreading. Without contractions, cells fail to spread to different areas on soft and rigid surfaces and to maintain monolayer integrity. We further observe that cadherin contractions enable cells to test myosin IIA-mediated tension of neighboring cells, and sort out myosin IIA-depleted cells. Thus, we suggest that epithelial cells test and respond to the mechanical characteristics of neighboring cells through cadherin contractions.

## Introduction

For the proper organization of tissues, cells need to probe the mechanical properties of their micro-environment including both extracellular matrix and neighboring cells through adhesive contacts. These mechanical properties are then transduced into biochemical information to regulate cell functions^1^, including single and collective cell motility^2, 3^, proliferation^4^ or differentiation^5^. Of the many mechanical properties that cells control, stiffness appears to be an important parameter that is distinctive for a tissue and is reflected in the cells that constitute the tissue^6^. It follows that cells should be able to measure the stiffness of their neighbors to enable them to regulate their cell-cell contacts, cytoskeletal rigidity and organize cell monolayers. Thus, it is important to understand how E-cadherin rigidity might be sensed. Recent studies have indeed found that epithelial cells spread to larger areas on rigid cadherin-coated surfaces than soft^7^. The testing of cadherin adhesion rigidity^8^ shares similarities with the testing of matrix rigidity described for fibroblasts^9^. In the context of epithelial cell dynamics, this mechanism may allow cells to adapt to changes in the local stiffness of their neighbors due to cytoskeleton remodeling and reinforcement^10–12^.

Cadherin rigidity is a complex mechanical parameter since it is defined as the force per unit area needed to displace a cadherin adhesion by a given distance. In the case of matrix rigidity sensing, cells pull matrix contacts to a constant deflection and measure the force generated^13–15^. The local matrix rigidity sensor is a sarcomere-like contraction complex (2 micrometers in length) that contracts matrix adhesions by 120 nm and if the force exceeds 25 pN, then a rigid-matrix signal is activated in the cell. The contractions are controlled by receptor tyrosine kinases in terms of the magnitude of deflection, the duration and the activation of the contractions^16, 17^. The sarcomere-like contraction system consists of antiparallel actin filaments anchored by α-actinin, a bipolar myosin filament and a number of actin binding proteins including tropomyosin 2.1^14, 15^. Although there are many obvious differences between cadherin and integrin adhesions^18^, a similar mechanism may be used to sense the rigidity of the E-cadherin contacts, i.e. neighboring cells.

Integrin and cadherin adhesions have many features in common, including an organization involving distinct nanometer-sized clusters of adhesion molecules^19, 20^ and many actin-binding proteins^18^. In tissues, cadherin clusters form homophilic interactions that maintain adhesions between cells^21^ and mechanically hold the tissue together. As primary components of adhesive contacts, cadherins are major parts of the mechanotransducing systems between cells^22, 23^, and are important for tissue morphology ^24^. Many cytoplasmic proteins link these adhesions to the cytoskeleton and provide mechanical continuity across the cell through a dynamic actomyosin network and other filamentous elements^25^. In addition, a “sarcomeric belt” structure was reported at apical cell-cell boundaries of epithelial cells, with non-muscle myosin II-mediated actomyosin structures interpolated in between cadherin clusters at a constant spacing^26, 27^. Other mechanical activities of epithelial monolayers also appear to involve actin and myosin contractions of the cadherin adhesions including the formation of cell-cell contacts^28^, the contraction and bending of cell monolayers^27, 29^ and tissue extension. The cadherin adhesion complexes are consequently a major element in mechanosensing events that ultimately shape the tissue and are involved in rigidity sensing and many other processes.

Previous studies have shown that cells generated high forces on large N-cadherin-coated pillars through cellular level contractions that were similar but not identical to matrix traction forces^8, 30^. N-cadherin-junctions that formed on N-cadherin-coated pillar surfaces resembled the morphology and dynamics of native epithelial cell–cell junctions^8^. Moreover, substrate stiffness modulated the level of force on E-cadherin adhesions that correlated with changes in cell spread area^7^. If the cadherin-based rigidity-sensing module was similar to the integrin-based sensor, then it should be evident in the deflection patterns of sub-micrometer diameter pillars^9^. When we placed E-cadherin expressing cells on sub-micrometer E-cadherin-pillars, we observed local contractile units of 1.2–2.4 micrometers that pulled E-cadherin junctions together to a constant distance of 130 nm, independent of rigidity over a 20-fold range. Unlike the integrin-based contractions, E-cadherin contractions did not require myosin IIA but were rather dependent upon myosin IIB, α-catenin, vinculin and also involved tropomyosin 2.1. When the contractions were depleted, monolayers did not form properly and there was improper sorting of mixed monolayers. The density of cadherin contractions correlated with the area of cells on cadherin surfaces, which was consistent with the increased spreading of MDCK cells on stiffer E-cadherin surfaces. Thus, it seems that cells create local contraction units between E-cadherin contacts to test mechanical properties of neighboring cells for proper organization of epithelial monolayers.

## Results

### COS-7 and MDCK cells form contractile units on E-cadherin-coated pillars

Since previous studies indicated that cadherin adhesion clusters are constantly spaced^28, 31^, especially that E-cadherin clusters were spaced at a distance of ~1.4 μm^28^. Based on these observations, we prepared pillar substrates with a diameter of 600 nm and 1.2μm center to center spacing. When pillars were coated with E-cadherin, cells attached and developed force on the pillars (Supp. Fig. 1A). To characterize E-cadherin-dependent force generation, we used COS-7 cells, an SV-40 transformed derivative of African Green Monkey Kidney Fibroblasts and Madin-Darby Canine Kidney (MDCK) epithelial cells. COS-7 cells were of particular interest because these fibroblast-like cells expressed E-cadherin^32^ while lacking a major myosin II isoform, Myosin IIA^33^, that was needed to produce local contractions on fibronectin matrices^17^. Surprisingly, COS-7 cells were able to adhere to, spread and pull on E-cadherin pillars and exhibited localized contractions (Fig. 1A, left panel). The spreading and force generation were similar to earlier studies using large cadherin-coated pillars^8, 34^; however, unlike the case with cells on the larger pillars, these sub-micrometer pillars revealed local contraction units of 1–2 micrometers like those previously found for integrin-based adhesions^9^. The criterion for the local contractions was that pairs of pillars moved toward each other for a limited period and maximum displacements occurred at approximately the same time point (see description below). When COS-7 cells spread on E-cadherin coated pillars, there were many examples of local contractions (Fig. 1A, right panel), which were not observed with larger pillars before, indicating that these smaller pillars were able to reveal local contractions in addition to the radial contractions.

**Figure 1.**
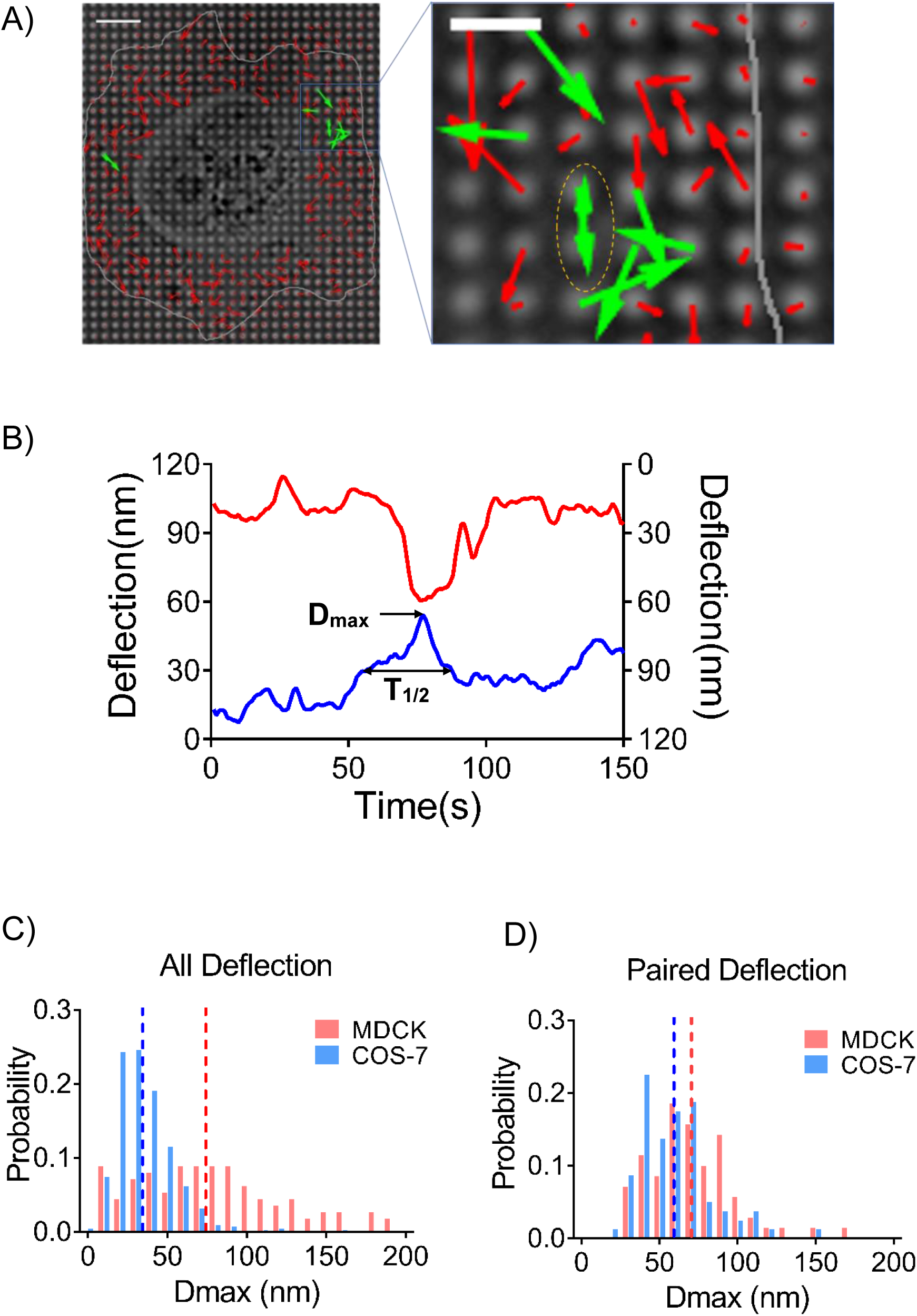
Cell generates local contractile units on E-cadherin coated pillars. A) Left: pillar deflection vector map under a COS-7 cell. Red vectors indicate non-paired deflections, green vectors indicate paired deflections, grey line indicates cell boundary. Scale bar 5μm. Right: Amplification of contraction-generation cell spreading area. B) Deflection plot of paired pillars under one contraction event, correlated with pillars indicated in A) in yellow dotted line circle. C) Histogram plots of total D_max_ distribution of MDCK cells was higher than that of COS-7 cells, dotted lines show mean D_max_ value of MDCK and COS-7 cells in respective color. D_max_=76.92 ± 4.332nm, n=112 for MDCK cells, and 35.7 ± 0.7837nm, n=661 for COS-7 cells. D) Histogram plots of D_max_ distribution of paired deflecting pillars in MDCK cells and COS-7 cells are similar. D_max_=71.1 ± 3.268nm, n=70 for MDCK cells, and 59.65 ± 2.678nm, n=80 for COS-7 cells.

In general, it was difficult to separate all local contractile units from radial contractions because multiple contractile units often overlapped resulting in complex pillar displacements. Although some local contractions were not recorded, we used the very stringent requirement for pairs that two pillars move toward each other and relax at the same time. The computer identification of pillar pairs involved two major criteria: 1) two pillars moved toward each other for more than 8 seconds during deflections of greater than half the D_max_, and 2) the D_max_ of both pillars occurred within 5 seconds. We then characterized paired contractions of pillars by two parameters: D_max_, the maximum pillar deflection value from the original position; and T_1/2_, half-peak contraction time that was the length of time that the pillar was pulled farther than half of the D_max_ value in a single pulling event (indicated in Fig. 1B). After analyzing the time course of pillar movements under spreading cells for ~30 minutes, there was a significant density of local contractile units, in which pillars deflected and relaxed in a synchronized manner (Fig. 1B, local contractions were noted by dotted line-circled pillars in Fig.1A). Characterization of the contractile units provided a quantitative analysis of the local contractions and we designated those paired E-cadherin adhesion-dependent contractile events as “**<u>c</u>**adherin **<u>c</u>**ontractions” (CC). It is important to note that the other pillar contractions that were not identified as pillar pairs had a much lower average displacement than the pillar pairs. Thus, the CCs were significant in density and were distinct from other pillar displacements.

To determine if CCs were present in other cells, we chose MDCK epithelial cells. After spreading on E-cadherin pillars for 3 hours, they generated cadherin contractions in a similar fashion to COS-7 cells (Supp. Fig. 1B). Analysis of the D_max_ of all pillar deflections showed that overall contractility was much greater with MDCK cells than with COS-7 cells because of the presence of Myosin IIA in MDCK cells (Fig. 1C), while the magnitude of CC deflection was similar in both cell lines (Fig. 1D). These results supported the idea that CCs were generated in similar fashion and were a common activity across different cell types in contrast to variations in overall force generation. Furthermore, analyses of the pillar deflections showed that the velocities of contraction and relaxation were equal in CCs (Supp. Fig. 1C); whereas the contraction velocities were significantly higher than relaxation velocities for total contractions in both MDCK and COS-7 cells (Supp. Fig. 1D). We also suggest that the CC pairs were unlike integrin-dependent contractions because they formed in the absence of Myosin IIA. In contrast, large contractions were much less frequent in overall deflections of COS-7 cells, indicating that large contractions observed in overall force in MDCK cells were indeed powered by Myosin IIA.

Since the local CCs were distinct and highly regular both in terms of D_max_ and duration of contractions, we quantified the CC parameters under a variety of conditions, including different pillar rigidities. When MDCK cells were spread on pillars with different rigidities due to their different heights, the local CCs had very similar D_max_ values (Supp. Fig. 2A, 71.1±27.3 nm on 0.75 μm high pillars, and 66.9±20.2 nm on 1.5 μm high pillars that had spring constants of 95 pN/nm and 12 pN/nm, respectively, data presented in mean±SD), indicating that contraction distance was rigidity-independent as was previously observed for local matrix contractions^14, 15^. Similar features were also observed for CCs generated by COS-7 cells spreading on E-cadherin pillars (D_max_ of 59.6±24.0 nm on 0.75 μm high pillars, and of 60.9±24.6 nm on 2 μm high pillars that had spring constants of 95 pN/nm and 5 pN/nm, data presented in mean±SD) (Supp. Fig. 2B). Further, the average T_1/2_ values were about 20.0 s for both COS-7 and MDCK cells (Supp. Fig. 3A). These results further reinforced the idea that paired contractions were powered by the same process in MDCK and COS-7 cells. Altogether, both MDCK and COS-7 cells produced CCs that were independent of rigidity and had similar D_max_ and T_1/2_ values. Thus, both MDCK and COS-7 cells pulled to a constant deflection and then the force of the contractions was proportional to the rigidity.

### E-cadherin-mediated rigidity response correlates with cadherin contraction density

In previous studies of matrix spreading, rigidity of the matrix was indicative of the density of matrix contractions and cell spread area^14, 17^. We then tested whether there was a similar correlation between CC density and spread area on E-cadherin-coated substrata. CC density was measured as the average number of CCs generated by a cell during 10 minutes in a constant area (36μm^2^ which is the area of 25 pillars). When MDCK cells spread on soft and stiff PDMS surfaces coated with E-cadherin, the final spread area was larger on stiff (~2 MPa) than on soft gel (~5 kPa) (Fig. 2A). As predicted from the local matrix contractions, the CC density was lower on the soft than on the rigid pillars (Fig. 2B), which correlated with the lower spread area on soft pillars (Fig. 2C). However, the density of COS-7 CCs increased on the soft pillars (Fig. 2D). This surprising result stimulated us to check the spreading of COS-7 cells on rigid and soft cadherin-coated surfaces. We observed that COS-7 cells spread to a larger area on soft (~5 pN/nm) than on rigid (~95 pN/nm) pillars (Fig. 2E), which correlated with higher CC density on soft pillars. These results indicated that CC formation was rigidity-sensitive and promoted cell spreading. However, COS-7 cells did not respond to rigid cadherin in the same way as MDCK cells. COS-7 cells exhibited transformed growth on soft fibronectin, which meant that they spread and grew equally well on soft and rigid fibronectin because they lacked rigidity-sensing contractions. Thus, the signal generated by CCs in COS-7 cells may have been different than in MDCK cells.

**Figure 2.**
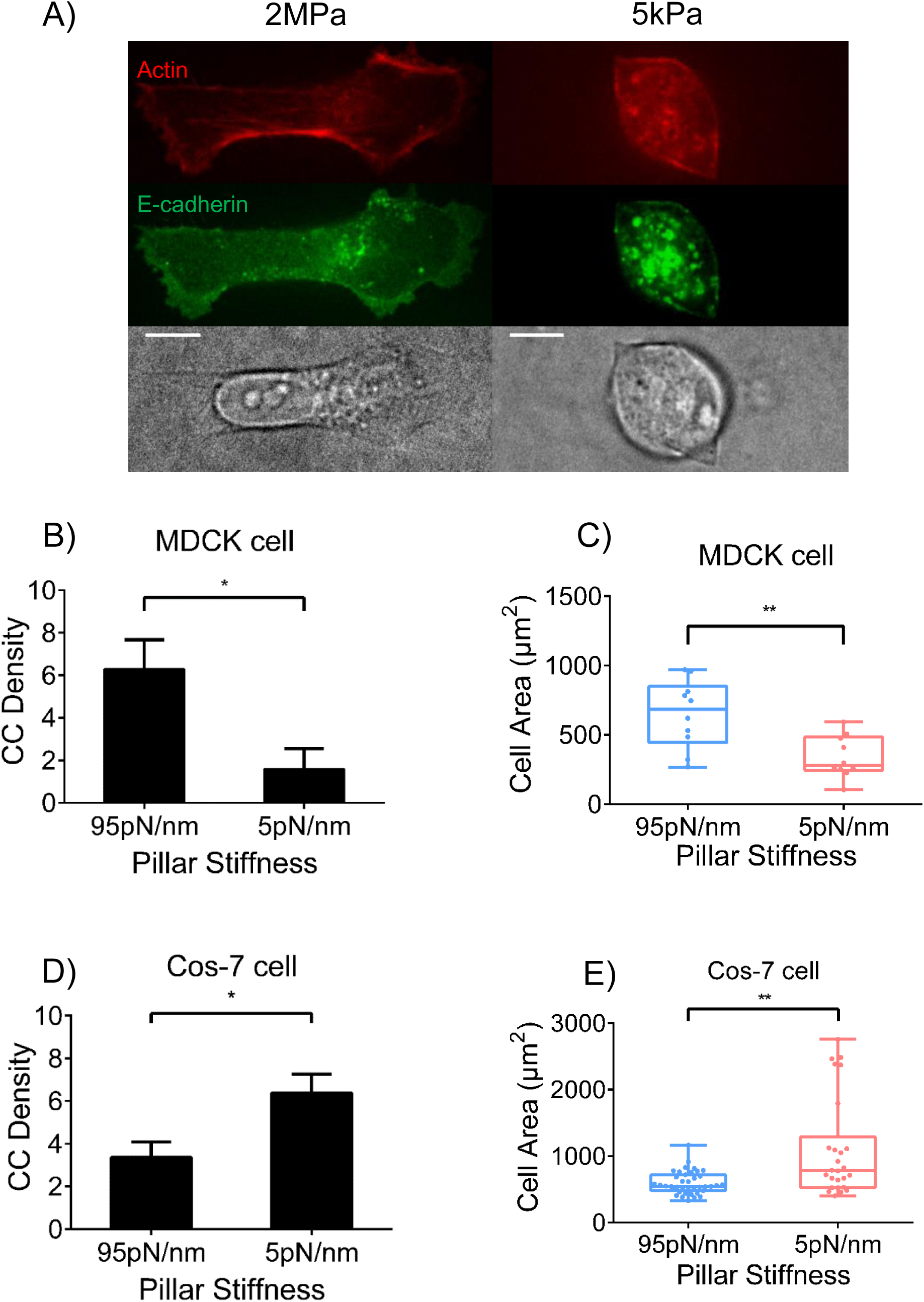
CC density correlates with cell spreading area on E-cadherin. A) On E-cadherin-coated PDMS gel, MDCK cells spread more on stiff 2MPa gel (left panels) than on soft 5kPa gel (right panels). Scale bar 10μm. B) CC density is higher when MDCK cells spread on stiffer (95pN/nm) pillars (6.29±1.39, n=5) than on softer (5pN/nm) pillars (1.60±0.95, n=4). C) MDCK cells spread more on stiffer pillars (650.1±78.1μm^2^, n=10) than on softer pillars (338.8±47.8μm^2^, n=10). D) CC density of COS-7 cells is higher on softer pillars (6.38±0.88, n=8) than on stiffer pillars (3.36±0.72, n=9). E) COS-7 cells spread more on softer pillars (1106.1±147.4μm^2^, n=26) than on stiffer pillars (598.1±27.5μm^2^, n=39). (*, p<0.05, **, p<0.01)

### α-catenin and vinculin co-operatively regulate cadherin contraction

To further investigate the role of cadherin adhesion proteins in CCs, we examined involvement of the major actin-binding proteins in the E-cadherin adhesions, α-catenin and vinculin. Previous studies showed that α-catenin was under force in the cadherin adhesion^35^, and perhaps acted as a molecular mechanosensitive switch^36^. There was also evidence for involvement of vinculin in linking adhesion complexes to actin^37^ that was consistent with its role as a force transducer. When MDCK cells stably missing α-catenin were placed on E-cadherin coated pillars, cadherin contraction density was greatly reduced (Fig. 3A). After α-catenin was restored in the knockdown cell line, a normal level of CCs was observed (Fig. 3B, paired CCs marked by green vectors) showing that α-catenin was critical in forming CCs (Fig. 3C). In addition, we also found that α-catenin knockdown reduced T_1/2_ value and D^max^ of the overall contractions, which was restored through α-catenin rescue (Supp. Fig. 3B-C). These results indicated that α-catenin was a crucial component in CCs and was generally involved in linking cadherin adhesions to the contractile cytoskeleton.

**Figure 3.**
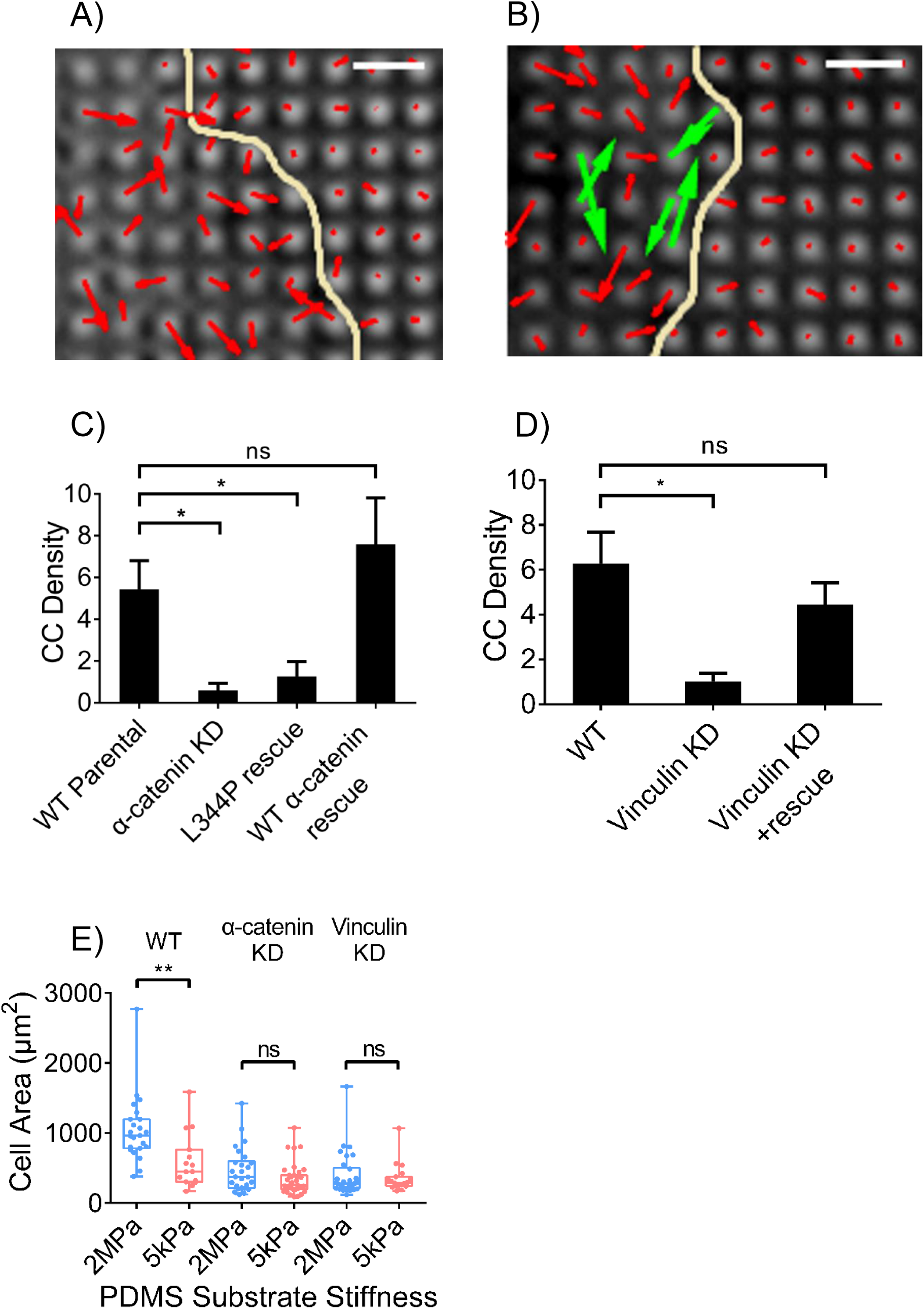
Depletion of α-catenin and vinculin in MDCK cells alters generation of CCs. Pillar deflection vector maps show that α-catenin KD MDCK cells fail to generate CC (A, red vectors indicate non-contractile deflections), while α-catenin rescue restores CC in KD cells (B, CC in green vectors). Yellow line indicates cell boundary. Scale bar 2μm. C) Quantification of CC density indicates that MDCK cells suffer from decreased CC density upon α-catenin KD (0.59±0.34, n=9), while L344P mutated α-catenin fails to restore CC (1.26±0.71, n=6), and wild-type α-catenin restores CC to normal density (7.59±2.23, n=4). D) Vinculin KD decrease CC density in MDCK cells (1.02±0.36, n=4), while vinculin rescue on KD background restores CC density (4.45±0.97, n=6). E) MDCK cells spread into larger areas on 2MPa E-cadherin coated gels (1049.4±99.9μm^2^, n=23) than on 5kPa gels (591.4±101.0μm^2^, n=15), while α-catenin KD diminishes such rigidity-dependence (464.8±61.8μm^2^ on 2MPa gel, n=26; 338.8±38.1μm^2^ on 5kPa gel, n=35), so is vinculin KD (392.6±57.1μm^2^ on 2MPa gel, n=30; 350.9±46.2μm^2^ on 5kPa gel, n=19). (*, p<0.05; **, p<0.01; ns, non-significant)

We next tested whether the association of vinculin with α-catenin was important. Although vinculin’s role in regulating force generation remained unclear, the interaction between vinculin and α-catenin depended upon the unfolding of α-catenin^36, 38^. To test if α-catenin-vinculin binding affinity affected cadherin contraction, we rescued α-catenin knockdown MDCK cells with an α-catenin mutant L344P that did not bind vinculin^39^. Upon L344P mutant rescue, local CC density was also significantly reduced compared with wild-type cells (Fig. 3C, Supp. Fig. 4A). Thus, the interaction between vinculin and α-catenin was important for CC formation.

To determine if vinculin was also involved in CC formation, we tested vinculin depleted MDCK cells and characterized their CCs on pillars. We found that vinculin knockdown cells had a much lower density of CCs when compared with wild-type MDCK cells, while re-expression of vinculin restored CC density to normal levels (Fig. 3D, Supp. Fig. 4B-C). We also measured the effect of vinculin depletion on magnitude and duration of overall contractions, but there was no significant difference in total contractions after vinculin knockdown (Supp. Fig. 4D-E). Thus, vinculin was involved in the CC mechanism even though vinculin depletion had no significant effect on overall cell contractility.

To further investigate the role of CC activity of MDCK cells in cell spreading, wild-type cells as well as α-catenin or vinculin depleted cells were spread on soft and rigid E-cadherin surfaces. We observed that wild-type MDCK cells spread more on rigid than on soft substrates, in agreement with previous results on pillars with varying stiffness (Fig. 2D, Fig. 3E). When cells lost ability to form cadherin contractions due to α-catenin or vinculin depletion, the cells spread similarly on soft and stiff substrates. In both cases, depleted cells spread less on the 2 MPa rigid substrate than wild-type cells (Fig. 3E). This was consistent with the hypothesis that CCs were involved in stabilizing the spread state and they required α-catenin and vinculin to form.

### Myosin IIB and Tpm2.1 mediate cadherin contraction

When MDCK cells spread on E-cadherin-coated pillar arrays, they were able to form individual E-cadherin clusters on pillar tips, but phosphorylated myosin light chain (pMLC) was found between E-cadherin clusters (Supp. Fig. 5A). This resembled fibroblast spreading on fibronectin-coated pillars in that integrins concentrated on pillars whereas pMLC was in between^9^. We also observed that treatment with Y-27632, a Rho-associated protein kinase (ROCK) inhibitor, fully abolished CC formation in COS-7 cells (Supp. Fig. 5B). These results indicated that myosin activity was critical for CC formation.

Previous studies indicated that all three types of non-muscle myosin II activity were involved in E-cadherin contact dynamics^27, 40^ and E-cadherin-based force generation^8^. However, COS-7 cells lacked non-muscle Myosin IIA and expressed primarily Myosin IIB plus a minor fraction of Myosin IIC^33^. Since the CC density was similar in COS-7 and MDCK cells, it was unlikely that Myosin IIA was involved in CCs. To determine whether Myosin IIB or Myosin IIC were involved in CCs, we immunostained COS-7 cells spread on E-cadherin substrata for Myosin IIB and Myosin IIC. Myosin IIB immunostaining co-localized with pMLC at the cell edge, where most of the CCs were found (Supp. Fig. 5C). In contrast, Myosin IIC did not localize with CCs in spread COS-7 cells and did not co-localize with pMLC (Supp. Fig. 5D). At a super-resolution level, phosphorylated myosin IIB bipolar minifilaments localized between two contracting pillars (Fig. 4A), suggesting direct involvement of Myosin IIB in CC formation. To confirm that myosin IIB was indispensable in CC formation, we knocked down myosin IIB with shRNA and introduced myosin IIA-GFP in COS-7 cells at the same time to create myosin IIA positive, IIB negative COS-7 cells (Fig. 4B). We found that these cells had many fewer CCs than did normal COS-7 cells (Fig. 4C), confirming that Myosin IIB, rather than IIA or IIC, was involved in CCs.

**Figure 4.**
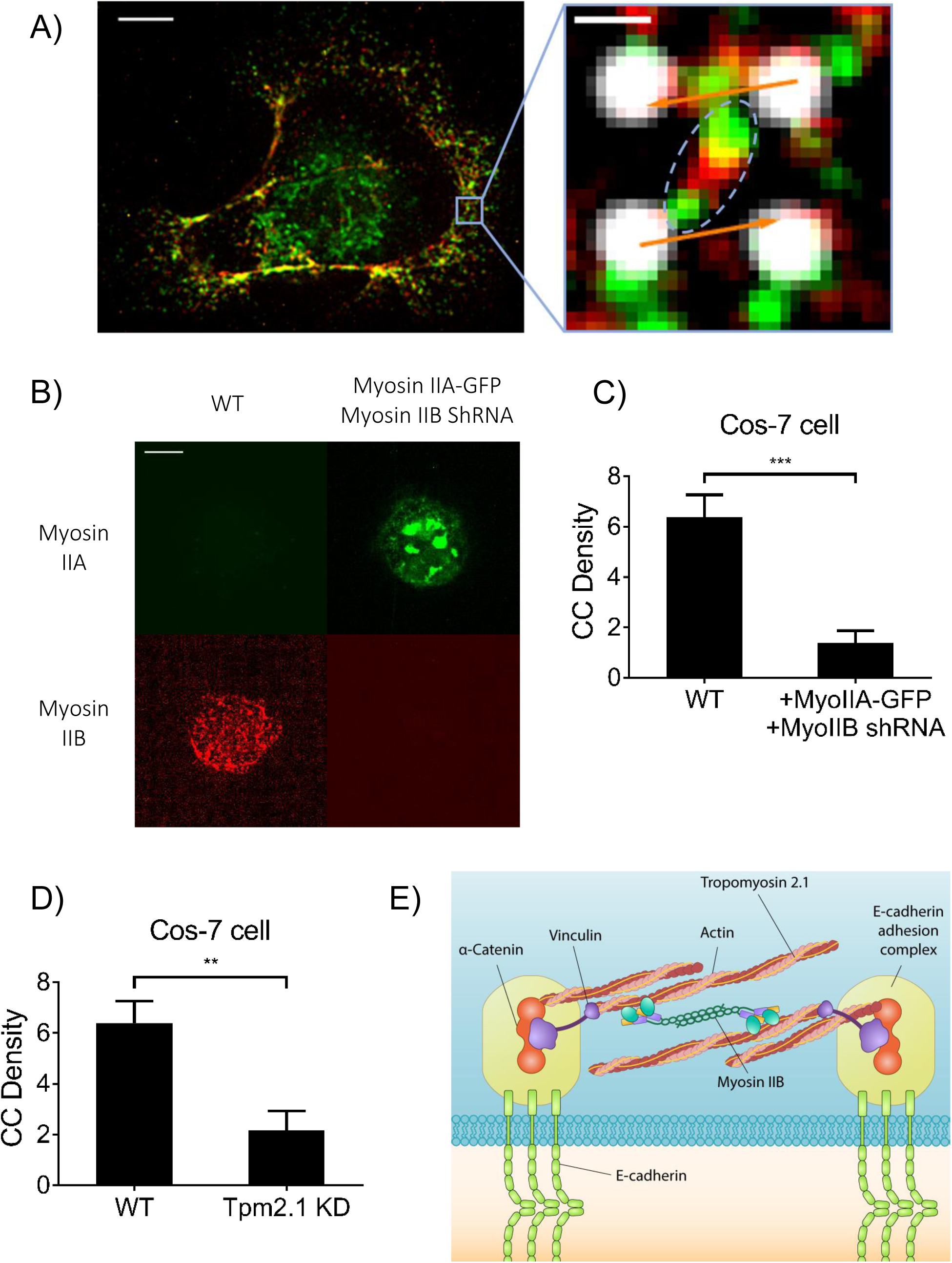
Non-muscle myosin IIB and Tpm2.1 mediates CC formation. A) Immunostaining of pMLC (in green) and myosin IIB heavy chain (in red) in COS-7 cell on E-cadherin pillars. Right panel shows phosphorylated myosin IIB mini-filament (shown in dotted line circle) between contracting pillars, deflections of which represented in orange arrows. Scale bar 5μm/0.5μm (left/right panel). B) Immunostaining of Myosin IIA and IIB in COS-7 cells (wild-type & with Myosin IIA-GFP and Myosin IIB shRNA). Scale bar 10μm. C) COS-7 cells expressing myosin IIA exhibited reduced CC density (1.37±0.75, n=5) upon myosin IIB knockdown compared to wild-type COS-7 cells (6.38±0.88, n=8). D) Tpm2.1 knockdown in COS-7 cells decreases CC density (2.18±0.75, n=6). E) Schematic representation of molecular mechanism of Cadherin Contraction. (**, p<0.01; ***, p<0.001)

Tropomyosin (Tpm) was identified as a major component in integrin contractile units, and cells were incapable of forming contractions or sensing rigidity when Tpm2.1 protein levels were downregulated^14^. Recent discoveries have indicated that several types of tropomyosin, notably Tpm2.1 and Tpm3, were involved in E-cadherin adhesion integrity^41, 42^. We found that Tpm2.1 accumulated in between E-cadherin-coated pillars in newly spread areas where CCs formed in the periphery of COS-7 cells, which resembled its localization in integrin contractile units ^14^. However, staining of Tpm3 showed that it was much more centrally located in the cells and did not overlap with CC-abundant areas or Tpm2.1 (Supp. Fig. 6A). We also found that pMLC localized to Tpm2.1-rich regions at the cell periphery (Supp. Fig. 6B). Moreover, we found that CCs localized at Tpm2.1-rich areas, where pMLC complexes were also observed to bridge between two contracting pillars (Supp. Fig. 6C). To confirm its involvement in CC formation, we knocked-down Tpm2.1 in COS-7 cells (see loss of Tpm2.1 staining pattern in KD cells, Supp. Fig. 6D). Tpm2.1 knockdown in COS-7 cells resulted in drastically reduction of CC formation (Fig. 4D), indicating that Tpm2.1 played an important role in CC assembly. Thus, we suggested that the CC unit between E-cadherin adhesions had a molecular basis distinct from local contractions of matrix (Fig. 4E).

### Alteration of Monolayer Organization with CC Depletion

Since both α-catenin and vinculin were indispensable in CC formation, we next tested the possibility that failure in CC formation would alter organization of cell monolayers. We seeded MDCK cells and let them grow to confluence for 2 days. 3D reconstruction of confocal images of actin showed that compared with the uniform monolayers formed by wild-type cells (Supp. Fig. 7A), α-catenin and vinculin knockdown cells formed abnormally organized monolayers, with protruding cells and less organized actin (Fig. 5A). 3D imaging of E-cadherin and actin organization in confluent monolayers showed that α-catenin knockdown induced formation of protruding cell aggregates above the basal monolayer, with actin and E-cadherin remaining localized at cell-cell boundaries (Fig. 5A, Supp. Fig. 7B). In comparison, vinculin knockdown cells had disorganized actin areas, and E-cadherin failed to properly localize at cell-cell boundaries as well (Fig. 5A, Supp. Fig. 7C). When we quantified the area of protruding cells above basal monolayer through analysis of actin staining distribution, we found that both α-catenin and vinculin knockdown cells had a significantly greater area protruding above cell monolayer than wild-type cells (Fig. 5B). We also performed a wound healing assay, and we observed that α-catenin knockdown caused a significant decrease in cell migration rate, and vinculin knockdown caused a mild decrease in migration rate (Fig. 5C, Supp. Fig. 7D). Of particular note, vinculin knockdown did not affect overall traction force on E-cadherin adhesions (Supp. Fig. 4E), but induced severe cell boundary disruption and collective migration retardation, which correlated with disruption of CC formation in the cadherin mechanotransduction process. These phenotypes correlated with depletion of CCs and indicated that CCs may have a role in maintaining normal epithelial integrity and collective cell migration.

**Figure 5.**
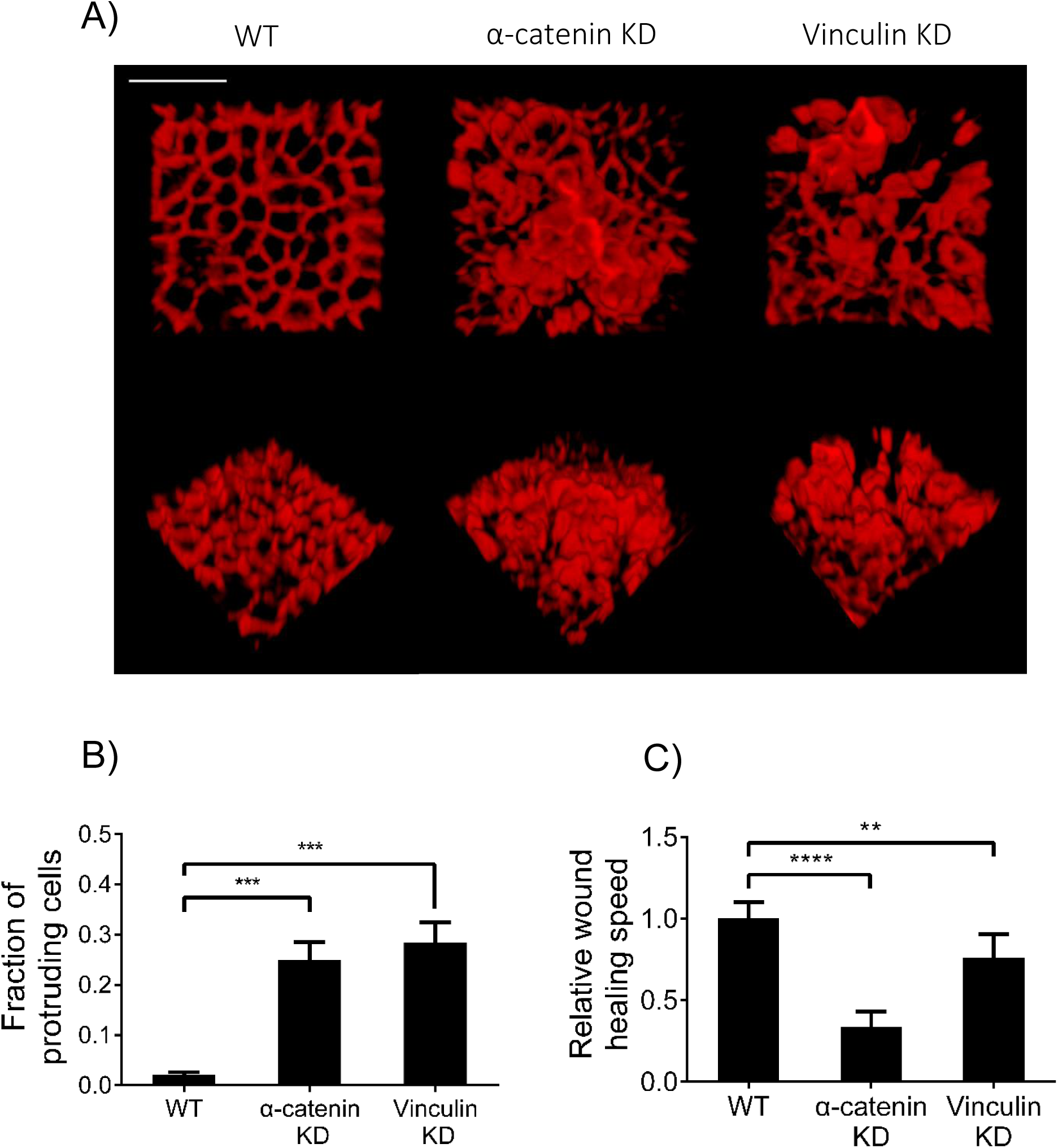
Deficiency of CC formation in MDCK cells induces abnormal monolayer formation and retarded collective migration. A) 3D reconstruction of actin (in red) in cell monolayers formed by wild-type (left panels), α-catenin KD (middle panels) or vinculin KD (right panels) MDCK cells. Upper panels show x-y views, lower panels show tilted 3D views. Scale bar 20μm. B) α-catenin KD or vinculin KD is able to induce higher protruding cell population in MDCK cell monolayers. (n=7 in each case, ***, p<0.001) B) α-catenin KD or vinculin KD decreases would healing speed in MDCK cell monolayers. (n=6 in each case, **, p<0.01; ****, p<0.0001)

### Cadherin Contraction regulates MDCK cell sorting in response to myosin IIA loss

Previous studies indicated that cell adhesion and cortical tension regulated cell sorting^43, 44^, and it was logical to propose that E-cadherin-mediated mechanotransduction at the cellular level was involved in this process. In order to test the role of CCs in cell sorting, we mixed cells with or without CCs and myosin IIA. Surprisingly, when MDCK and COS-7 cells (lacking myosin IIA) were mixed and co-cultured, the mixed cells exhibited clear segregation with MDCK cells pushing out COS-7 cells into islands surrounded by MDCK cells. In contrast, CC-deficient α-catenin knockdown MDCK cells commonly mingled with COS-7 cells (Supp. Fig. 8A). Since COS-7 cells lacked endogenous expression of myosin IIA, it appeared that myosin IIB-driven CC units to tested myosin IIA-mediated tension of neighboring cells. Through knocking-down myosin in MDCK cells, we observed that myosin IIB knockdown significantly disrupted CC level compared with wild-type or myosin IIA-KD MDCK cells (Supp. Fig. 8B). Wild-type MDCK cells identified myosin IIA-deficient COS-7 cells as aberrant cells despite their ability to generate CC and responded to the lack of cell contractility by segregating COS-7 cells. The ability to segregate COS-7 was dependent upon the presence of CCs, since α-catenin and vinculin KD MDCK cells that were unable to form CCs or sense E-cadherin rigidity were also unable to segregate COS-7 cells and randomly mingled with them. Thus, it appeared that CCs were involved in organizing epithelial monolayers.

To test the generality of this hypothesis, we mixed MDCK cells with or without CCs and myosin IIA knockdown MDCK cells. Mingling was quantified from the segregation area of myosin IIA positive cells. We found that all wild-type MDCK cells sorted themselves from myosin IIA knockdown cells and formed large areas of isolated knockdown cell islands. CC-deficient MDCK cells, caused by depletion of either α-catenin, vinculin, or myosin IIB, produced much smaller cell islands, and showed greater mingling with myosin IIA-KD MDCK cells (Fig. 6A, quantified in 6B). We also quantified the time-dependence of the segregation using the segregated cell area. Wild-type cells produced a much greater segregated cell area of the myosin IIA-KD cells over 24 to 48 hours post-mixing, while CC-deficient cells (α-catenin-KD cells), showed no change in the level of segregation from myosin IIA KD cells between 24 and 48 hours (Supp. Fig. 8C, quantified in Fig. 6C). These results indicated that CCs were sensing the level of myosin IIA in their neighbors and that sensing was necessary for the segregation of myosin IIA deficient cells (Fig. 6D-E).

**Figure 6.**
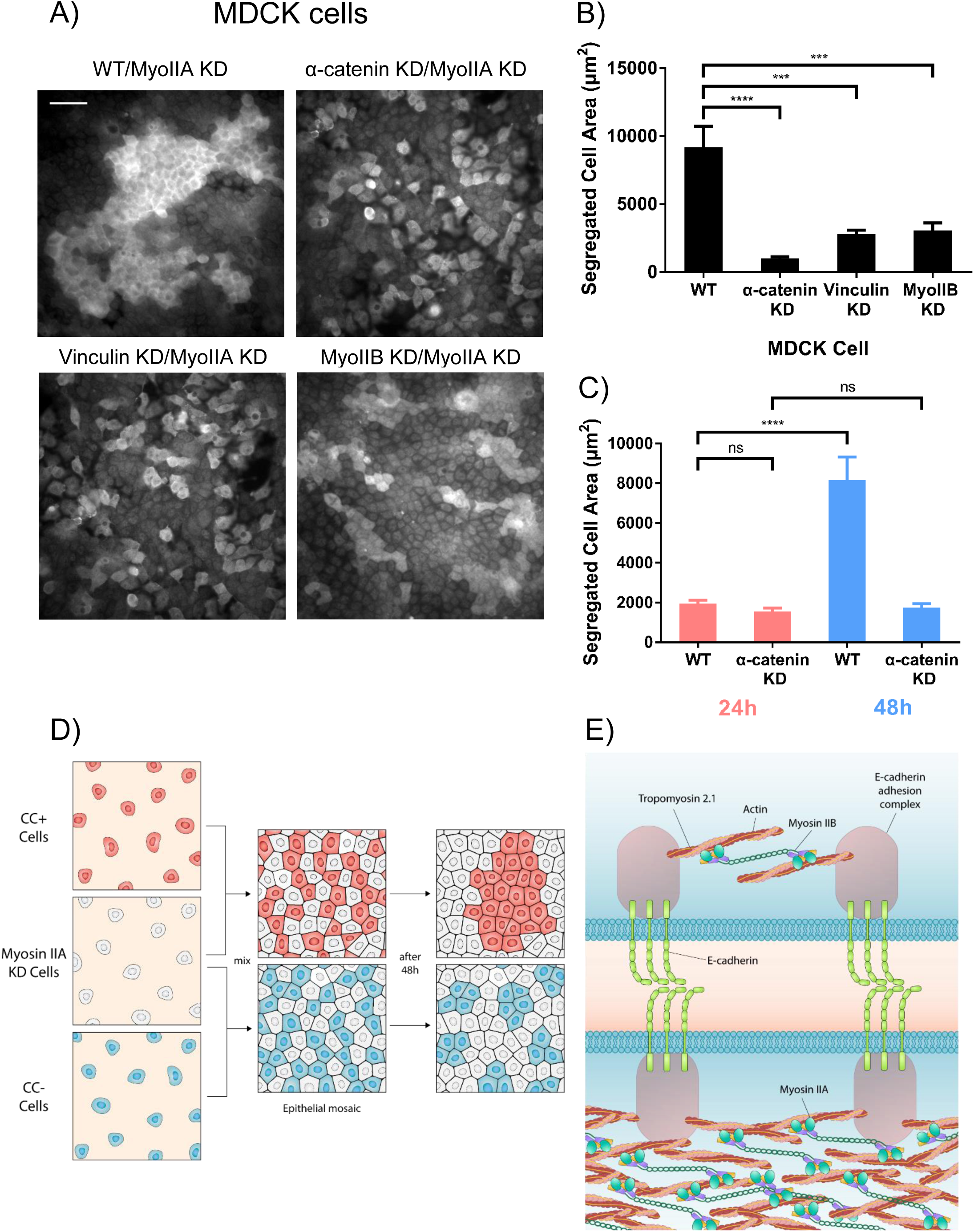
Cadherin contraction regulates MDCK cell sorting. A) Myosin IIA immunofluorescence indicates populations of MDCK cells (wild-type, α-catenin-KD, vinculin-KD, Myosin IIB-KD) mixed with Myosin IIA-KD MDCK cells. Scale bar 50μm. B) Quantification of isolated MDCK cell areas after mixed with Myosin IIA-KD MDCK cells for 48 hours (wild-type, n=30; α-catenin-KD, n=50; vinculin-KD, n=60; Myosin IIB-KD, n=50). C) Quantification of isolated MDCK cell (wild-type or α-catenin-KD) areas after mixed with Myosin IIA-KD MDCK cells for 24 (wild-type, n=65; α-catenin-KD, n=50) or 48 hours (wild-type, n=32; α-catenin-KD, n=55). D) Schematic representation of CC-regulated, myosin IIA-sensitive cell sorting. “CC+ cells” refers to cells capable of forming CCs, “CC-cells” refers to cells unable to form CCs. E) Schematic representation of cell-cell rigidity sensing, Myosin IIB-mediated CC tested myosin IIA-mediated tension across neighboring cell boundaries. (**, p<0.01; ***, p<0.001; ****, p<0.0001)

## Discussion

In this study, we find that paired contractions of E-cadherin coated pillars correlate with the ability of the cells to sense cadherin rigidity and to form epithelial monolayers. During CCs, pillar pairs are contracted by a total of 120–140 nm for a period of about 20 s irrespective of pillar rigidity over a nineteen-fold range of rigidity (from ~5 to ~95 kPa). As predicted from previous studies of the proteins in cadherin junctions, α-catenin, vinculin, myosin IIB and tropomyosin 2.1 were needed for the CC units to form (Fig. 4E). Of the cadherin adhesion complex proteins, α-catenin and vinculin could anchor actin filaments to the adhesions. The CCs are distinct from local fibronectin matrix contractions in that they do not rely upon Myosin IIA but rather require Myosin IIB as well as tropomyosin 2.1. Further, they are distinct from overall radial contractions of E-cadherin-coated pillars in length of deflection and velocity. Although CC pairs form preferentially in newly spread areas of the cell, they continue after cells are spread and are involved in maintenance of cell-cell adhesions. Also, we find CC density to be rigidity sensitive since density increases with increasing rigidity in MDCK cells and decreases with increasing rigidity in COS-7 cells. Further, CC density changes correlate with changes in spread area on different rigidity cadherin surfaces. The organization of epithelial monolayers is highly dependent upon the CCs for both the morphology of the monolayer and the sorting of cells in the monolayer. Thus, it seems that the CCs are important for the formation and maintenance of normal epithelia.

Local contractions between E-cadherin adhesions provide a simple mechanism for testing the rigidity of neighboring cells that is analogous to local matrix contractions to test matrix rigidity even though the details are distinct. Many physiological processes are postulated to involve cadherin adhesion mechanosensing such as convergent extension ^45^ and epithelial tissue movements^46^. In those cases, changes in monolayer morphology are coupled with continued cell-cell sensing, and this indicates that there is a general mechanism of E-cadherin sensing in tissues^6^. Recent studies indicate that the rigidity of cadherin-coated surfaces affects cellular responses, indicating that the cells can sense cadherin rigidity^7, 8^; however, little is known regarding how cells employ force-generating molecular complexes to test rigidity through cadherin adhesions. The presence of the local contractions provides a simple mechanism for probing cell rigidity, because pulling cells contract to a constant distance and sense the force, which can be a simple measure of the rigidity of the neighboring cell. In a general context, although there is only a poor understanding of how contractile force is converted into a signal for rigidity, the many similarities between matrix rigidity sensing and cadherin rigidity sensing make it logical to propose that analogous mechanisms are involved.

In the case of the cell-matrix rigidity sensing, the complex is similar to a sarcomere in terms of the components and the organization. Basically, anti-parallel actin filaments cover the 1.5–2.5 micrometer gap between matrix adhesions and myosin II bipolar filaments contract them at a very slow rate of 2–3 nm/s. Similarly, the CCs are driven by myosin II between two adhesions and the velocity of contractions is 2–3 nm/s. The major differences are in the type of myosin II with Myosin IIB powering CCs and Myosin IIA powering matrix contractions and in the primary cadherin complex proteins, α-catenin and vinculin, that anchor the actin filaments needed for CCs.

In terms of the mechanism of linkage with the actin cytoskeleton, the CCs depend strongly upon α-catenin and vinculin. Both proteins are involved in actin binding to the E-cadherin complex^10^. Knockdown of α-catenin reduces the number of contraction events. This indicates that α-catenin serves as an important mechanical linker between cadherin adhesions and actomyosin. In addition, α-catenin is generally an important component for force transmission at the cellular level. Similarly, depletion of vinculin causes a dramatic decrease in the density of CCs. Further, decreasing the vinculin binding affinity of α-catenin also reduces CCs. If vinculin binding is important for stable actin linkages to α-catenin^35, 47^, then it is not surprising that alterations that weaken vinculin binding to α-catenin inhibit CC formation.

The roles of α-catenin and vinculin in CCs extends to their regulatory role in tissue integrity. α-catenin has been identified as a tumor suppressor, and its depletion could trigger YAP-1 mediated overgrowth^48^. Similarly, disruption of myosin-powered contractility also induces YAP-associated contact inhibition failure^49^. We find that either α-catenin or vinculin knockdown causes disorder in cell monolayers. As expected, depletion of vinculin significantly disrupts actin organization and E-cadherin localization at cell-cell boundaries concomitant with the loss of CCs; however, depletion does not alter overall force generation behavior of cells in both magnitude and duration (Supp. Fig. 4D-E). Thus, the effects of either α-catenin or vinculin knockdown clearly indicate that they have indispensable roles in establishing the epithelial monolayers perhaps through aspects of cadherin contractions.

Involvement of Tpm2.1 again highlights that cadherin contractions and integrin contractions share key molecular components. As an important component in both rigidity sensing processes, the role of Tpm2.1 in CC formation provides another insight into rigidity insensitivity of Tpm-deficient cancer cells^14, 50^. Cancer cells are insensitive to matrix rigidity, and without CCs would also not be able to test and respond properly to the rigidity of neighboring cells in a tissue. Previous studies also show that Tpm2.1 knockdown in epithelial cells retards wound closure^41^, which agrees with our wound healing assay results in CC-deficient MDCK cells with α-catenin or vinculin knockdown. These results further support a significant role for Tpm2.1 in regulation of tissue integrity and cancer suppression through cadherin mechanotransduction, in addition to its role in cell-matrix rigidity sensing^14^.

In cases where all of the cells in a monolayer lack the CCs, there is improper organization of the monolayer. Many cells lacking CCs lose adhesion to the glass substrate and leave the surface in irregular folds of the monolayer. This indicates that the CCs are an important part of the development of the proper boundary morphology and the columnar nature of the cells in a monolayer. COS-7 cells will form normal cell-cell adhesions and do not overgrow in monolayers (unpublished data). However, the monolayers are very flat with a large area per cells because of the lack of myosin IIA to create tension needed to form columnar cells in a monolayer. Mechanosensing through CCs appears to contribute to the proper sorting of cell-cell contacts in monolayers and we suggest that the signaling from the CCs to the myosin IIA contractile network is a critical step. Thus, mechanosensing appears to be an important aspect of monolayer organization.

The traditional theory to explain cell sorting attributes cell segregation to cell adhesion-mediated tension, and the sorting process then minimizes free energy and boundary length of different cell populations^51^. How cell-cell adhesion contributes to the sorting is largely unexplained. These observations indicate that an asymmetrical process occurs at cell-cell boundaries, namely CCs sense myosin IIA-mediated tension in neighboring cells. If myosin IIA is missing, normal cells will sort themselves from myosin IIA-null cells. But if the cells lack CCs and cannot test their neighbors, they co-mingle with cells lacking myosin IIA. In terms of the relation to surface rigidity, the CCs on rigid pillars will develop stronger contacts than on soft and hence will cause the cell to spread more on rigid pillars. Spreading of the myosin IIA-null COS-7 cells on cadherin pillars could be the result of the transformed nature of those cells or a result of the loss of linkage between the sensing system and myosin IIA contractile networks. Thus, cells with CCs can be recognized as “abnormal” cells if they lack of myosin IIA, whereas CC-deficient cells, if expressing normal levels of myosin IIA, will be recognized as “normal” cells and escape from being sorted out. Such a mechanosensing system can identify weak or dying cells and can help to explain part of the normal sorting behavior.

The transient nature of the E-cadherin contraction units is consistent with the transient nature of many cellular mechanosensing processes^52^. In the MDCK cells, the transient contractions cause a response of more contractions on rigid pillars than on soft pillars, which correlates with greater spread area. Although many possible factors could contribute to CC density, we find more CCs in COS-7 cells on soft than on rigid pillars and the cells spread more on soft. This can logically relate to the absence of myosin IIA in COS-7 cells, since CCs are testing the myosin IIA contractility. In cell sorting assays, cells with low myosin IIA levels may naturally sort away from those with high levels because of the downstream responses. The molecular mechanisms of signaling need to be worked out in these systems to understand the phenomena.

In the case of integrin contractile units, rigid matrices produce high forces that cause increased tyrosine phosphorylation and increased adhesion strength. During spreading on matrices, the initial tests help to reinforce adhesions and the rate of testing drops over the first hour. In the case of CCs, the rate of testing doesn’t appear to change dramatically with time, implying that cells are continually checking on their neighbors’ cortical tension. If the neighboring cells lose intracellular tension or provide improper mechanical feedback, the sorting reactions could be initiated, including possible expulsion of the low-tension cells. Further, CCs are different in cells with different physiological backgrounds, indicating that they have a ubiquitous role in probing the surrounding cells. Thus, characterization of cadherin contractions can provide insight into E-cadherin mediated mechanosensory events, enabling a better understanding of how cells interact with their neighbors to create a proper epithelial monolayer.

## Materials and methods

### Cell lines and culture

All MDCK cell lines (ATCC) and COS-7 cells (ATCC) were cultured in high-glucose, L-glutamine containing DMEM with 10% of fetal bovine serum (FBS). For all assays using E-cadherin coated surfaces, high-glucose, L-glutamine containing DMEM without FBS, and supplemented with 100U/mL penicillin/100μg/mL streptomycin was used in experiments. All reagents were from Thermofisher. MDCK cell lines (GFP- and mCherry-tagged E-cadherin) were acquired from W. J. Nelson’s lab^53^. Stable knockdown of α-catenin in MDCK cells was performed with shRNA in W. J. Nelson lab^54^. The Vinculin knockdown MDCK cell line was from Dr. Soichiro Yamada’s group in University of California, Davis^55^. Myosin IIA KD and myosin IIB KD MDCK cells were from René-Marc Mège Lab. COS-7 cell line was from Michael Sheetz lab.

### Plasmids and transfection

Non-muscle myosin IIB shRNA and vinculin-GFP plasmid was a gift from Dr. Alexander Bershadsky lab in MBI, vinculin-GFP originated from Michael Davidson lab in Florida State University. α-Catenin plasmids (wild-type and L344P mutant) and GFP-E-cadherin plasmids were described earlier^39^. Tpm2.1 knockdown was performed with siRNA oligonucleotides as previously published^14^.

Neon® Transfection System (Thermo Fisher) was used for electroporation of MDCK cells. Lipofectamine 2000 (Thermo Fisher) was used for chemical transfection of COS-7 cells.

### Preparations of nanopillar arrays and flat gel substrates

We used sub-micron size pillars for recording and analyzing cell contraction behavior. The pillars were in a square pattern, 600nm in diameter (D), and with three different heights (L) of 750nm, 1500nm and 2000nm. Pillars were made of PDMS (Sylgard® 184 Silicone Elastomer Kit), mixed at a ratio of 10:1, spin-coated on silicon molds and cured at 80°C for 2 hours. For pillars of 600nm in diameter, bending stiffness was calculated to be ~95nN/μm for 750nm tall pillars, ~12nN/μm for 1500nm tall pillars, and ~5nN/μm for 2000nm tall pillars, actual stiffnesses were quantified as previously published^9^. Pillars were patterned in a square grid, with neighboring centroid-to-centroid distance of 1.2μm (2D) or 2.4μm (4D). Flat PDMS gel surfaces were prepared on glass coverslips with Sylgard 184 silicone elastomer kit (Dow Corning). PDMS surfaces with a Young’s modulus of 2MPa were made with elastomer to curing agent ratio of 10:1, and 5kPa surfaces were made with ratio of 75:1, as previously published ^56^.

For E-cadherin coating, PDMS films with polymerized pillars were peeled from silicon molds, placed on 12 mm glass bottom dishes (Iwaki) and treated with O_2_ plasma for 5 minutes. Then, they were incubated with 10μg/ml anti-human Fc antibody (Jackson research, goat anti human) in 0.1M borate buffer (pH=8) at 4°C overnight. For flat PDMS substrates, samples were treated by the same procedure without surface plasma treatment. Coated substrates were washed with DPBS three times, and reacted with 10μg/ml E-cadherin-Fc chimera protein (R&D systems, diluted in DPBS containing Mg^2+^ and Ca^2+^) for 2 hours in room temperature, and washed with DPBS three times before use.

### Cell spreading assay, wound-healing assay, drug treatment and immunostaining

MDCK cells (wild-type and all knockdown lines) were trypsinized and replated onto E-cadherin coated pillar arrays at low density in serum-free media as mentioned above, and incubated at 37 °C for 3 hours for cells to properly adhere to pillars before transfer to the microscope for imaging. For spreading assays on PDMS gels, cells were replated onto E-cadherin coated PDMS gels at low density in serum-free media and incubated at 37 °C for 6 hours before fixation and staining. COS-7 cells were trypsinized and replated onto E-cadherin coated pillars and imaged immediately since they started to spread in a rapid manner. For wound-healing assays, cells were cultured to near-confluent density on 35mm Petri Dishes (Nunc), and then scratched in the central area of the confluent cell monolayers. Cells were then cultured in Biostation IMQ microscope (Nikon) for long-term imaging.

For Y-27632 (Y0503, Sigma) treatment, COS-7 cells were treated with 10μM Y-27632 for 2 hours, resuspended with drug-containing serum-free media and seeded onto pillar substrates for image acquisition.

For cell immunostaining in general, cells were fixed with 3.7% paraformaldehyde (PFA, diluted in DPBS containing Mg^2+^ and Ca^2+^) for 15 minutes at 37 °C, permeabilized with 0.1% Triton X-100 for 10 minutes at room temperature, and blocked with 1% bovine serum albumin (BSA)/DPBS solution (blocking buffer) for 1 hour before staining with primary antibody in blocking buffer at 4 °C overnight. Samples were washed four times with DPBS before secondary antibody staining in blocking buffer for 1 hour at room temperature and washed four times afterwards before DAPI staining for 5 minutes. Phalloidin staining was applied together with secondary antibody incubation for 1 hour.

Primary antibodies used in these experiments are listed as following: Phospho-Myosin Light Chain 2 (Ser19, Mouse mAb #3675 and Rabbit mAb #3671, CST), non-muscle myosin IIA (M8064, Sigma), non-muscle myosin IIB (CMII 23, DSHB), non-muscle myosin IIC heavy chain (PRB-444P, Covance), Tpm2.1 (TM311, Sigma), Tpm3.1/3.2 (ab180813, Abcam), E-cadherin (610181, BD Biosciences).

### Cell Sorting Assay

For MDCK cell-COS-7 cell mixture, MDCK cells (WT or α-catenin KD) and COS-7 cells were trypsinized, counted and mixed at a 1:1 ratio, and seeded at a total density of 5.6×10^5^ cells per dish after centrifugation and re-suspension. For MDCK cell mixture, myosin IIA-positive MDCK cells (WT, α-catenin KD, vinculin KD or myosin IIB KD) were mixed with myosin IIA KD MDCK cells at 1:4 ratio, and seeded at a total density of 7×10^5^ cells per dish. Mixed cells were incubated in 12mm glass-bottomed dish (IWAKI) for 48 hours before immunostaining unless otherwise specified.

Myosin IIA was stained as mentioned above in all mixed monolayers to distinguish between MDCK cells and COS-7 cells, or between myosin IIA positive and negative MDCK cells. Cell areas with positive myosin IIA immunostaining intensity were thresholded and quantified to indicate the level of sorting segregation between different types of cells.

### Microscopy imaging and data analysis

Cell spreading on pillars was imaged with a DeltaVision system attached to an Olympus IX71 inverted microscope with x100 oil immersion objective (1.40NA, UPlanSApo) and Photometrics CoolSNAP HQ2 (CCD) camera. SoftWoRx (4.10) software was used to control the imaging configuration and recording. Fluorescence images of cells on pillars or PDMS gels were acquired using a spinning-disc confocal microscope (PerkinElmer) attached to an Olympus IX81 inverted microscope. Super-resolution confocal images were acquired using Live-SR (Roper Scientific) module attached to a Nikon Eclipse Ti-E inverted Microscope body, controlled by MetaMorph (7.10.1.161), iLas (1.2.0) and Live-SR (1.7.3) software. Biostation IMQ (Nikon) was used to record the wound healing process, cell samples were incubated with 5% CO_2_ and were imaged with 10x objective for up to 24 hours.

Pillar position detection was conducted in ImageJ (NIH) through tracking plugins designed by Dr. Felix Margadant. Pillar deflection analysis was conducted through Matlab (Math works). Statistical analysis and graph plotting were generated through Prism (Graphpad software), all bar plots were presented as Mean±SEM. All data were presented in Mean±SEM unless otherwise specified. Cadherin contraction detection program was adapted from Matlab codes used in previous studies of integrin contractile units^17^. Analyses of significant difference levels were performed using unpaired Student’s t-test with Welch’s correction.

## Supporting information

## Acknowledgements

We thank all the members of the Sheetz and Mège-Ladoux labs for their help, especially F. Margadant, S. Liu and B. Yang for their help in designing ImageJ plugins and Matlab codes for pillar analysis. We thank the Core Facilities of Mechanobiology Institute. Y. Yang and E. Nguyen are supported by Mechanobiology Institute, National University of Singapore and Ministry of Education, Singapore. R.-M. Mège is supported by Fondation ARC, Projet International de Coopération Scientifique (PICS) CNRS programmes, France-Singapour exchange grant, National University of Singapore-Université Sorbonne-Paris-Cité (NUS-SPC) exchange grant and Fondation pour la Recherche Médicale (FRM). B. Ladoux is supported by Mechanobiology Institute, the European Research Council under the European Union’s Seventh Framework Program (FP7/2007-2013) / ERC grant agreements n° 617233, Agence Nationale de la Recherche (ANR) “POLCAM” (ANR-17-CE13-0013) and USPC-NUS program. M. P. Sheetz is supported by NIH grants, NUS grants and Mechanobiology Institute, National University of Singapore.

## Author Contributions

Y.Y. and E.N. performed the experiments; S.N.G.H.N. and M. H. prepared the myosin KD MDCK cell lines; Y.Y. analyzed the data; Y.Y., B.L. and M.P.S. designed the study; Y.Y., R.-M.M., B.L. and M.P.S. wrote and prepared the manuscript.

