## Supplementary material for "Local Contractions Regulate E-Cadherin Adhesions, Rigidity Sensing and Epithelial Cell Sorting"

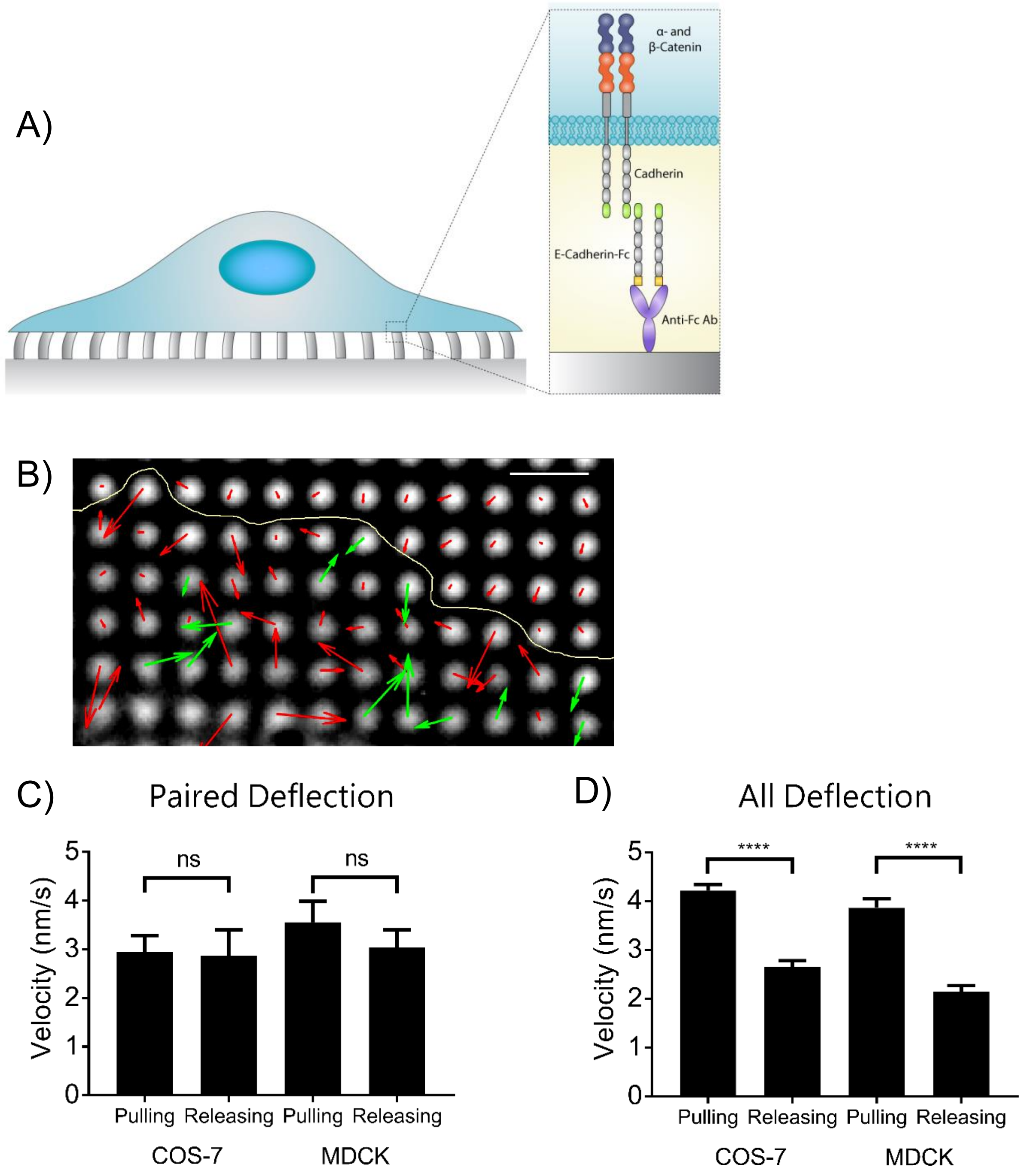


Supplementary Figure 1. Cadherin contraction in MDCK cells and velocity analysis of pillar deflection. A) Schematic representation of cell adhering on E-cadherin coated pillars. B) Vector map of pillar deflections under one MDCK cell. Red vectors indicate non-paired deflections, green vectors indicate paired deflections. Scale bar 2µm. C) CC pulled and released pillars in similar speeds in both COS-7 (pulling velocity=2.95±0.34nm/s, n=16; releasing velocity=2.87±0.53nm/s, n=16) and MDCK cells (pulling velocity=3.56 ±0.43nm/s, n=16; releasing velocity=3.04±0.36nm/s, n=16). D) Unspecific pillar deflections exhibited significant difference between pulling and releasing speeds in COS-7 (pulling velocity=4.21±0.13nm/s, n=402; releasing velocity=2.65±0.13nm/s, n=402) and MDCK cells (pulling velocity=3.87±0.19nm/s, n=195; releasing velocity=2.14±0.13nm/s, n=195) . (****, $p<0.0001$; ns, non-significant)

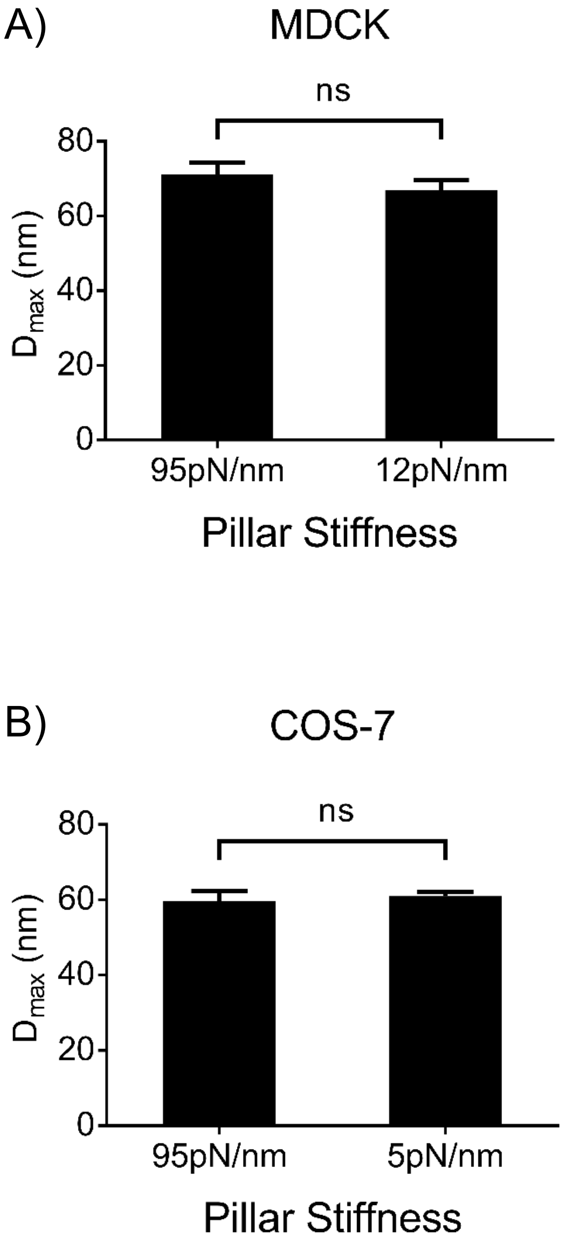


Supplementary Figure 2. Substrate stiffnesses regulates CC density, but not contraction length. A) Bar plots of $D_{max}$ of CC generated by MDCK cell on pillars with stiffnesses of 95 ($D_{max}$=71.1±3.27nm, n=70) or 12pN/nm ($D_{max}$=66.92±2.81nm, n=52). B) Bar plots of $D_{max}$ of generated by COS-7 cell on pillars with stiffnesses of 95 ($D_{max}$=59.65±2.68nm, n=80) or 5pN/nm ($D_{max}$=60.94±1.19nm, n=428). (ns, non-significant)

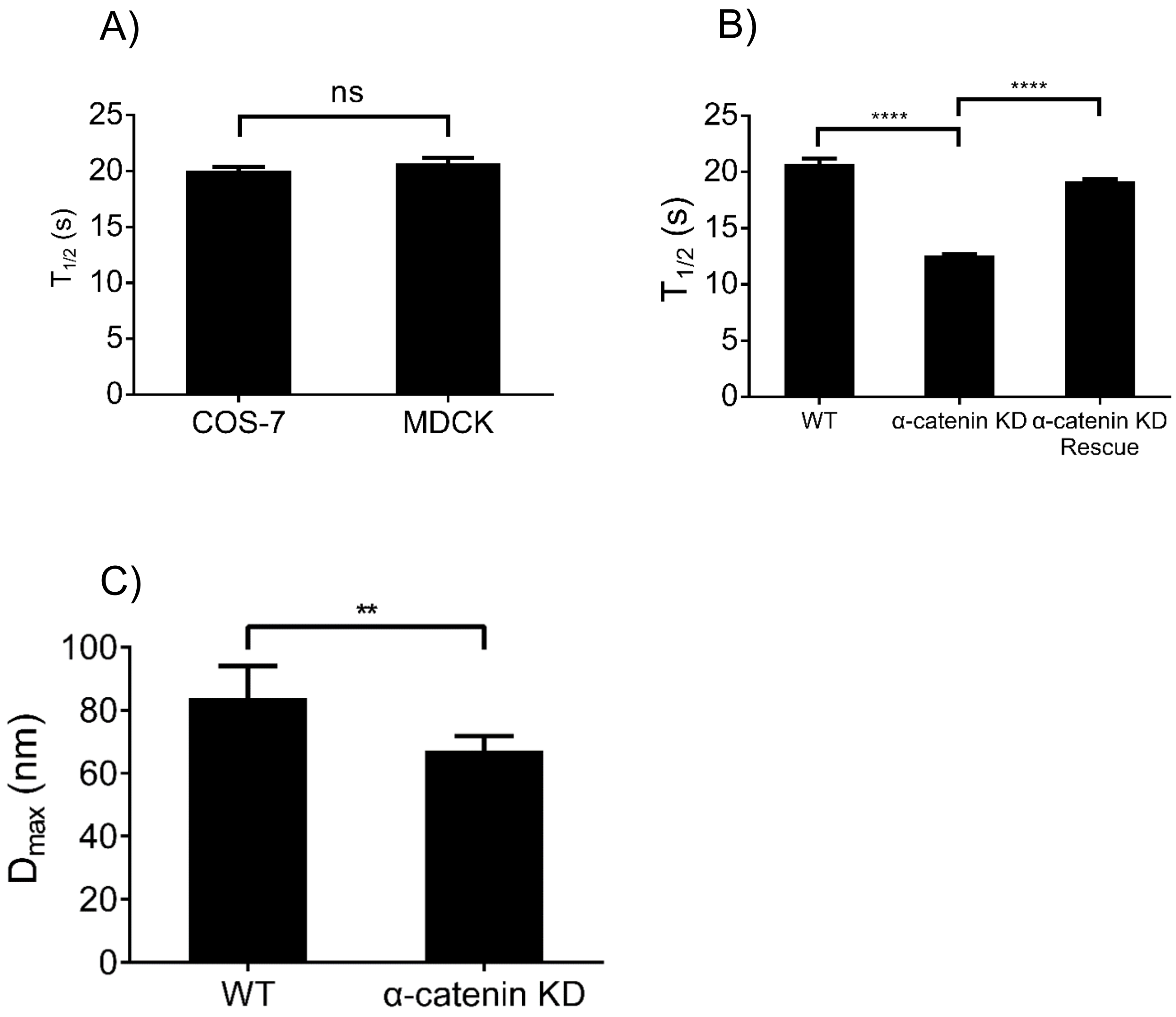


Supplementary Figure 3. Time duration of E-cadherin mediated force generation. A) Bar plots of pulling half-peak time distributions of COS-7 cells (20.08±0.33s, n=438) and wild-type MDCK cells (20.69±0.53s, n=157) on pillars. B) α-catenin knockdown significantly reduced average pulling half-peak time of MDCK cells, and rescue α-catenin recovered pulling half-peak time to normal level (WT, $T_{1/2}$=20.69±0.53s, n=157; α-catenin knockdown, $T_{1/2}$=12.55±0.14s, n=2789; α-catenin rescue, $T_{1/2}$=19.20±0.17s, n=2789). C) α-catenin knockdown significantly reduced overall $D_{max}$ distribution in MDCK cells (WT, $D_{max}$ =83.77±5.14nm, n=117; α-catenin knockdown, $D_{max}$=67.27±2.30nm, n=275). (**, $p<0.01$; ****, $p<0.0001$; ns, non-significant)

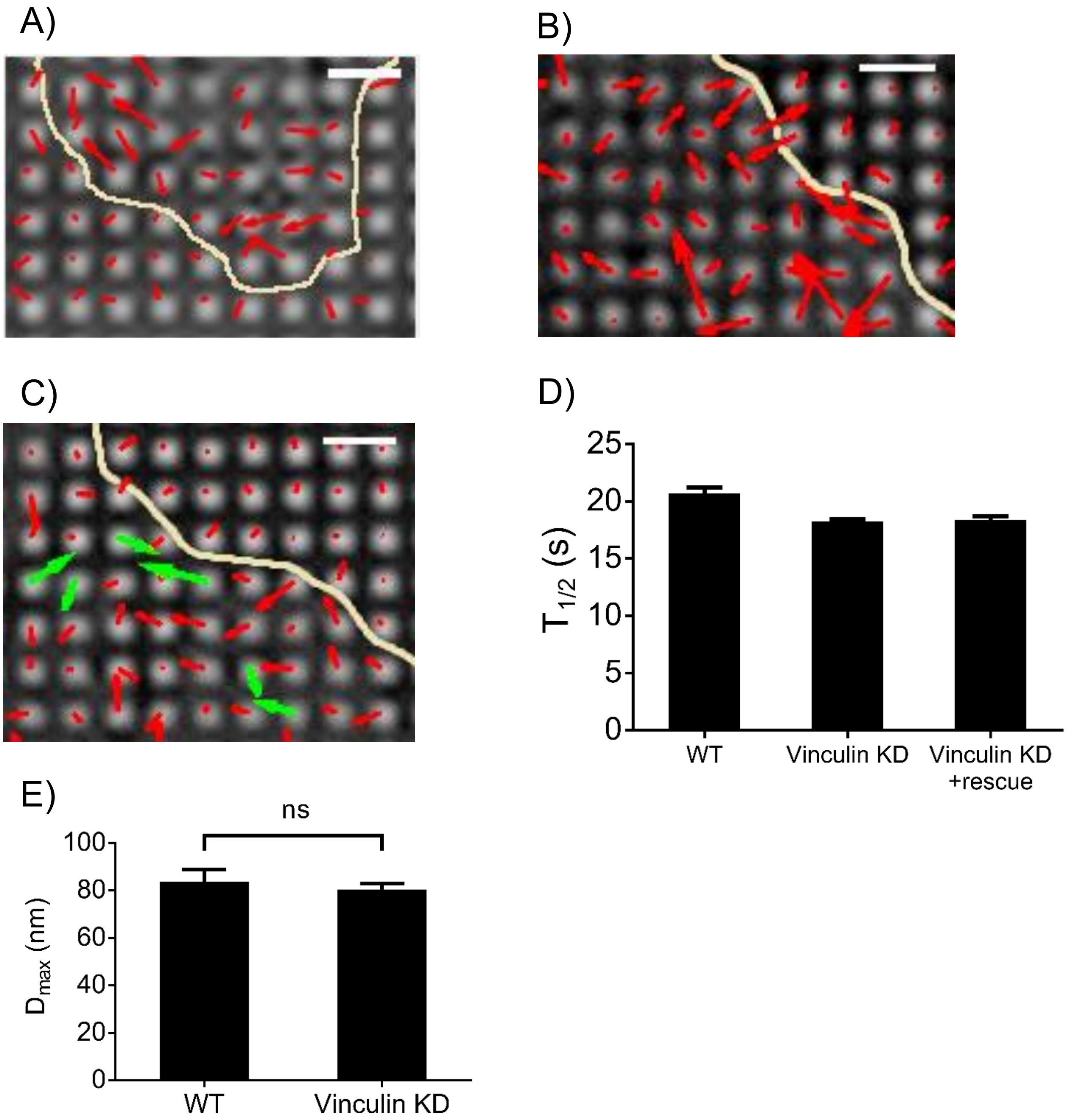


Supplementary Figure 4. Vinculin depletion impairs cadherin contraction, but not overall orce generation. Vector map showed that A) α-catenin KD rescued with L334P and B) vinculin KD MDCK cells failed to generate CCs at normal level, and C) vinculin rescued under KD background restored CCs. Vectors in red indicate non-paired pillar deflection, those in green indicates paired deflection. Yellow line indicates cell boundary. Scale bar 2μm. D) Vinculin knockdown did not significantly change average half-peak time of MDCK cells on pillars (WT, $T_{1/2}$=20.69±0.53s, n=157; vinculin knockdown, $T_{1/2}$=18.27±0.19s, n=1988; vinculin rescue, $T_{1/2}$=18.42±0.30s, n=572). E) Vinculin knockdown did not alter overall $D_{max}$ distribution in MDCK cells (WT, $D_{max}$ =83.77±5.14nm, n=117; vinculin knockdown, $D_{max}$=80.42±2.48nm, n=200). (ns, non-significant)

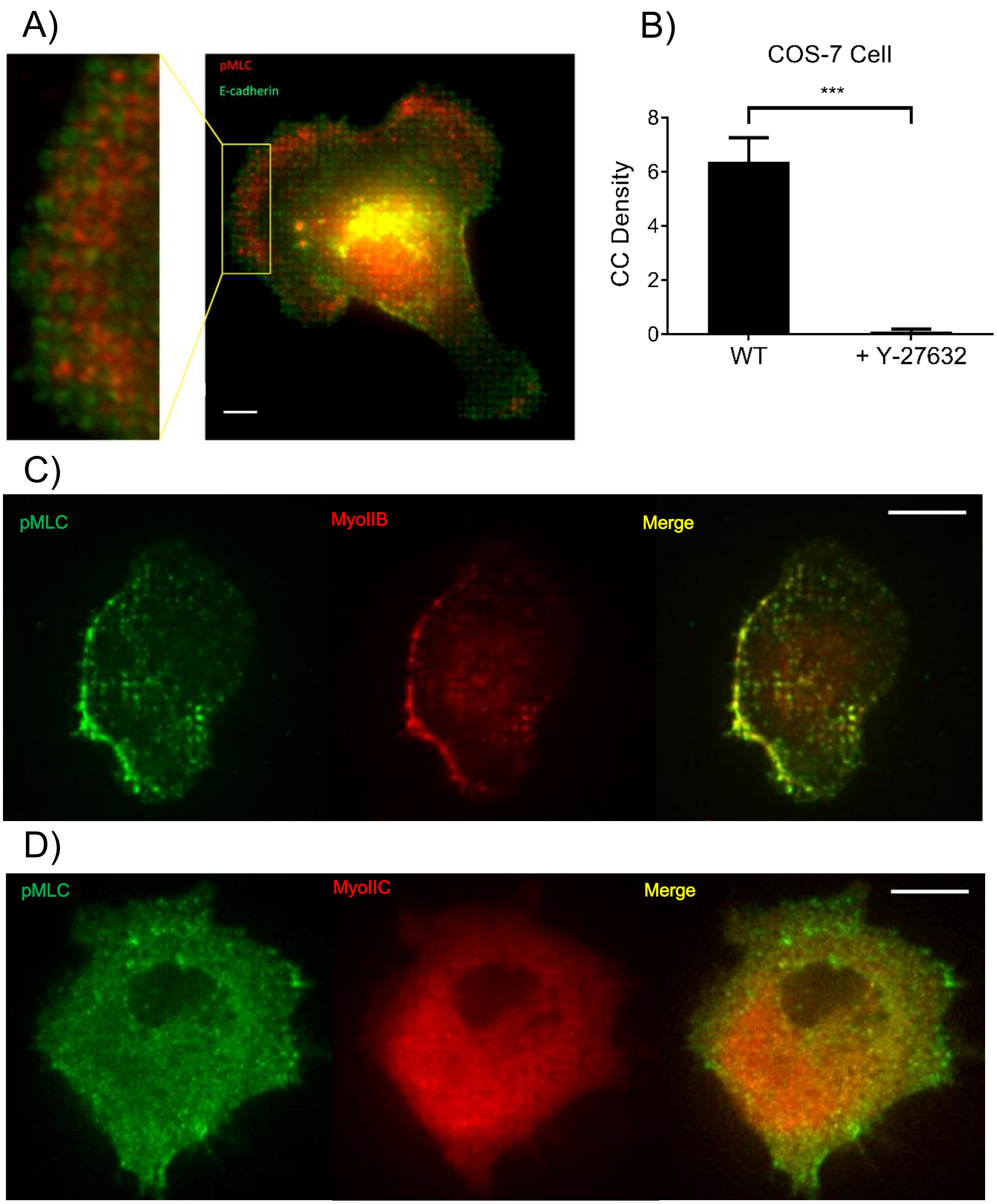


Supplementary Figure 5. Myosin phosphorylation is required for cadherin contraction. A) Phospho-myosin light chain localized between individual E-cadherin adhesion on the pillar tips at periphery area of an MDCK cells. Scale bar 5µm. (n=5 in each case, * **p<0.001) B) Y-27632 treatment (10uM for 2 hours) decreased CC density in COS-7 cells. C-D) Co-localization of Myosin IIB/IIC and pMLC in Cos7 cells spreading on E-cadherin pillars. Left panels show pMLC localization, middle panel in C) shows localization of myosin IIB, middle panel in D) shows localization of myosin IIC. Right panels show merge of both channels. Scale bar 10µm.

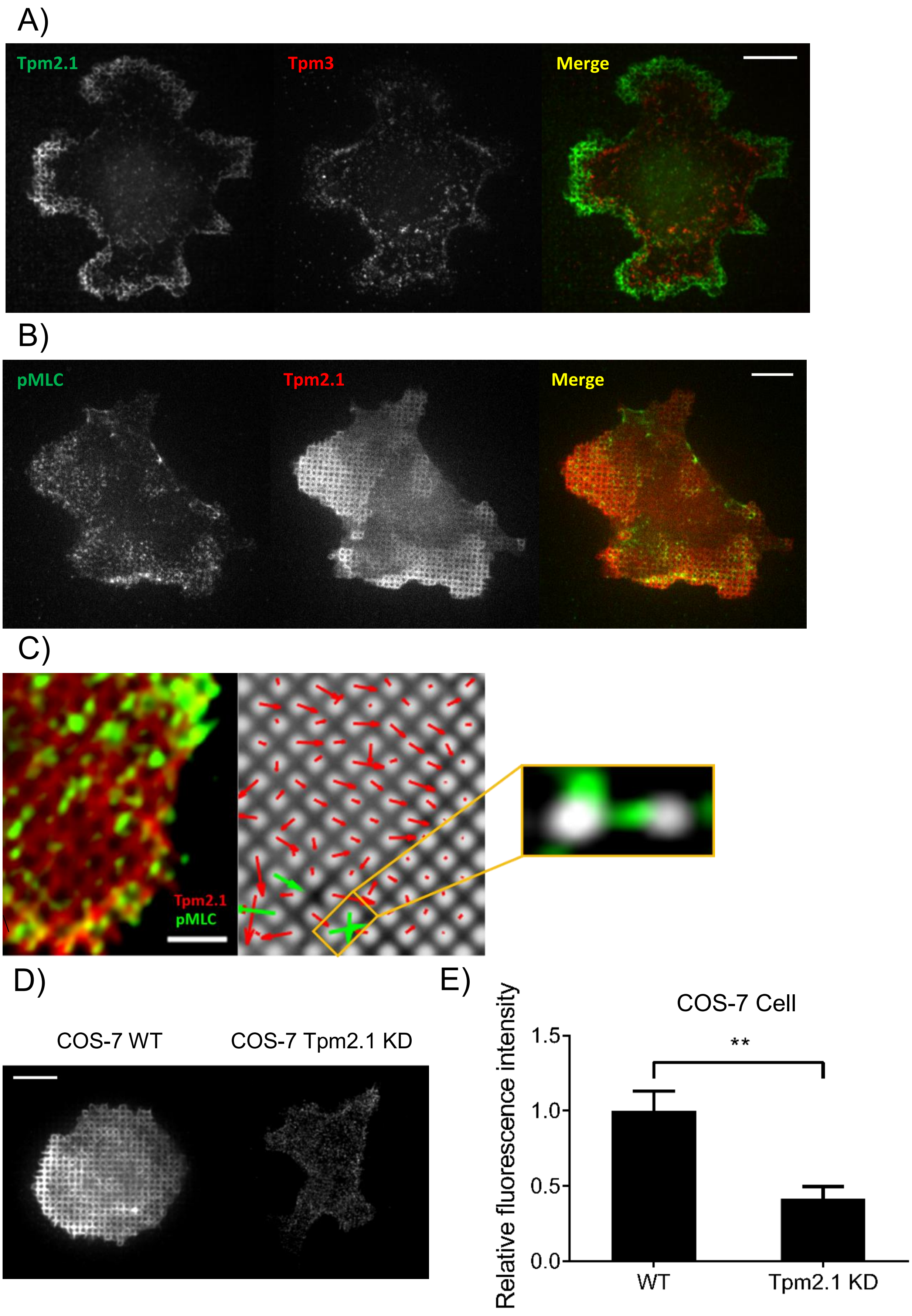

A)
Tpm2.1
Tpm3
Merge
B)
pMLC
Tpm2.1
Merge
C)
Tpm2.1
pMLC
D)
COS-7 WT
COS-7 Tpm2.1 KD
E)
COS-7 Cell
Relative fluorescence intensity
1.5
1.0
0.5
0.0
**
WT
Tpm2.1 KD

Supplementary Figure 6. Localization of tropomyosin in COS-7 cells on E-cadherin pillars. A) Localization of Tpm2.1 and Tpm3.1/3.2 in COS-7 cells on E-cadherin coated pillars. Scale bar 12μm. B) Immunostaining of pMLC and Tpm2.1 in COS-7 cells on E-cadherin coated pillars. Scale bar 10μm. C) Super-resolution image of Tpm2.1 (red) and pMLC (green) in CC unit. Left panel shows merged image, middle panel shows pillar deflection vector map at the fixation timepoint, red arrows indicate non-paired deflections, and green arrows indicate paired deflections. Right panel shows pMLC localized in contracting pillars (indicated in orange box in middle panel). Scale bar 2μm. D) T311 antibody staining of wild-type (left panel) and Tpm2.1 knockdown (right panel) COS-7 cells. Scale bar 10μm. D) Bar plots of relative T311 staining intensity of wildtype and Tpm2.1 knockdown COS-7 cells. (**, $p<0.01$)

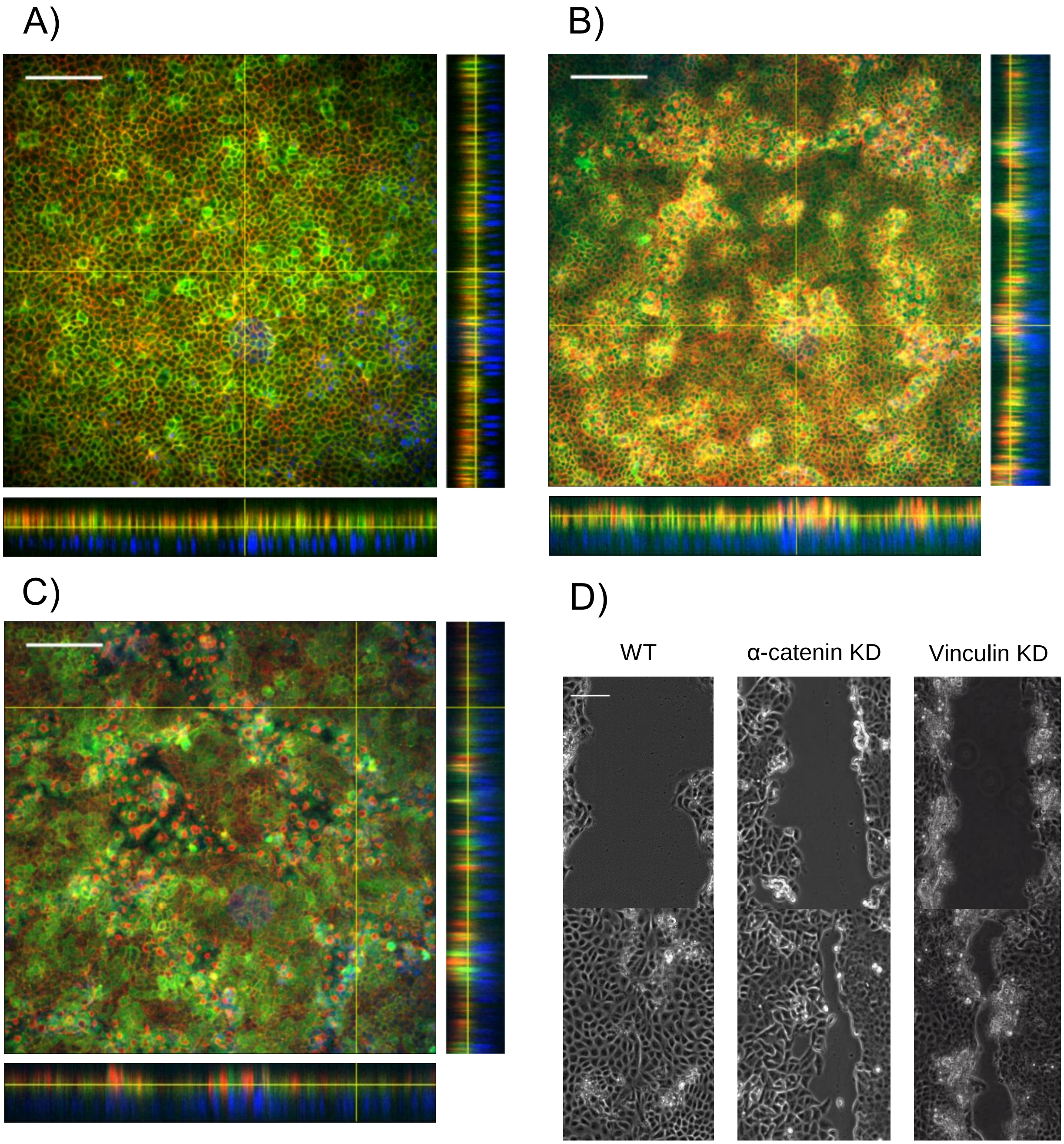


Supplementary Figure 7. Knockdown of α-catenin and vinculin disrupts epithelial organization and wound healing of MDCK cell monolayers. Z-stack images of A) Wild-type, B) α-catenin knockdown and C) vinculin knockdown MDCK cell layers are shown in orthogonal view. (Red: actin; Green: E-cadherin; Blue: DAPI). Scale bar 100µm. D) MDCK cell monolayers were imaged immediate after wounded (upper panels) and after 10 hours of wound closure (lower panels). Scale bar 100µm.

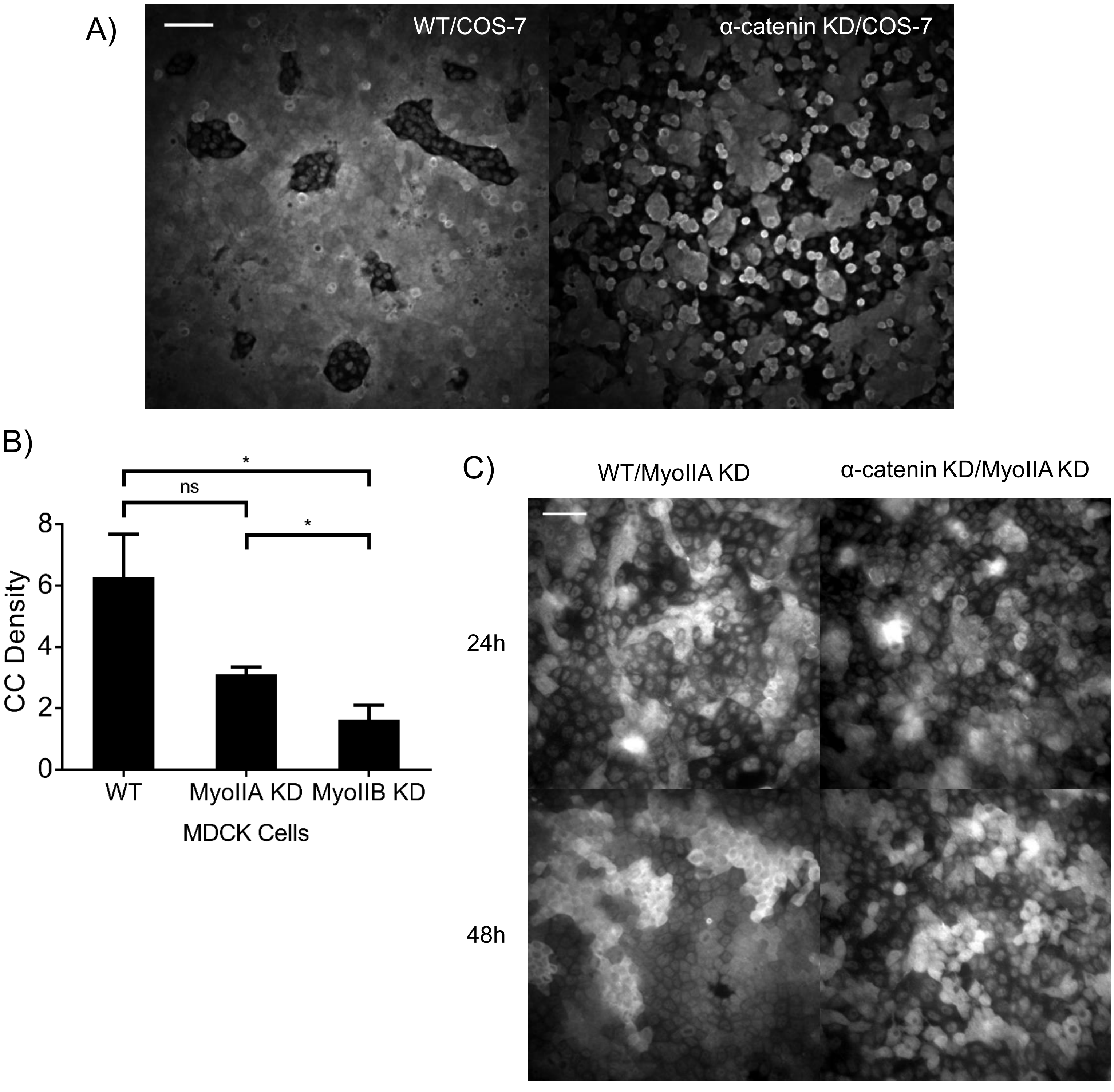


Supplementary Figure 8. CC regulates cell sorting through identifying myosin IIA. A) Myosin IIA immunofluorescence indicates populations of MDCK cells (wild-type and α-catenin-KD) mixed with COS-7 cells for 48 hours. Scale bar 100μm. B) Myosin IIA knockdown result in mild decrease of CC generation (3.12 ± 0.24, n=6) compared with wildtype MDCK cells (6.29±1.39, n=5), while myosin IIB knockdown results in significant disruption of CC generation compared with wildtype and myosin IIA-KD cells (1.64 ±0.47, n=6). C) Myosin IIA immunofluorescence indicates populations of MDCK cells (wild-type and α-catenin-KD) mixed with myosin IIA knockdown MDCK cells for 24 and 48 hours. Scale bar 50μm.
